# Mechanochemically Programmed, Oligomer-Selective Amyloid Assembly via Axial Rotation

**DOI:** 10.64898/2026.05.25.727554

**Authors:** Seokbeom Roh, Eunji Song, Minwoo Bae, Da Yeon Cheong, Sechan Han, Taeha Lee, Dain Kang, Hyungbeen Lee, Insu Park, Gyudo Lee

**Affiliations:** Department of Biotechnology and Bioinformatics, Korea University, Sejong 30019, Republic of Korea; Interdisciplinary Graduate Program for Artificial Intelligence Smart Convergence Technology, Korea University, Sejong 30019, Republic of Korea; Digital Healthcare Center, Sejong Institute for Business and Technology (SIBT), Korea University, Sejong 30019, Republic of Korea; Department of Digital Healthcare Engineering, Korea University, Sejong 30019, Republic of Korea; Department of Biomedical Engineering, Yonsei University, Wonju 26493, Republic of Korea

**Keywords:** Amyloid aggregation pathway, Amyloid oligomers, Mechanochemical control, Hydrodynamic boundary control, Cytotoxicity

## Abstract

Amyloid oligomers have been widely implicated as primary cytotoxic intermediates; however, their selective and scalable production remains challenging due to rapid fibril amplification. Here, we demonstrated that sustained axial rotation enables programmable mechanochemical control over amyloid pathway selection without the use of chemical additives. Using a thermal axial rotator, native monomeric hen egg-white lysozyme was incubated at 60 °C under quantitatively tunable rotational speeds, imposing defined centrifugal forces and wall-associated shear that restructured the hydrodynamic boundary conditions. A discrete transition emerged near 600 RPM, separating the two distinct assembly regimes. Below this threshold, aggregation followed a fibril-amplifying pathway characterized by elevated β-sheet content and elongated fibrillar morphologies. Above this threshold, fibrillar growth was strongly attenuated and oligomer-dominant assemblies predominated. Spectroscopic analyses and atomic force microscopy revealed that axial rotation redistributes the amyloid assembly states rather than simply suppressing aggregation. Functionally, the RPM-defined assemblies exhibited kinetically distinct seeding behaviors and induced divergent cytotoxic phenotypes in SH-SY5Y neuroblastoma cells. These findings establish axial rotation–mediated hydrodynamic boundary control as a scalable, chemical-free strategy for reprogramming amyloid assembly pathways and producing oligomer-rich assemblies with well-defined structural and functional properties.

## 1. INTRODUCTION

Amyloid protein aggregation—the conversion of soluble proteins or peptides into β-sheet–rich supramolecular assemblies—is implicated in a broad range of human diseases, including Alzheimer’s and Parkinson’s diseases, type II diabetes, and systemic amyloidosis [1–3]. Although the deposition of fibrillar amyloid aggregates as insoluble plaques or inclusions constitutes a pathological hallmark of disease in affected tissues, increasing evidence indicates that mature fibrils are not necessarily the principal cytotoxic species [2, 4–6]. Instead, smaller, soluble oligomeric intermediates formed along the aggregation pathway are increasingly recognized as the major drivers of cellular dysfunction and disease progression [5–7].

Despite their pathological relevance, amyloid oligomers are intrinsically difficult to isolate and characterize as discrete, well-defined species. Their low abundance, structural heterogeneity, and metastable nature promote continuous interconversion with monomers and progressive maturation into fibrils, complicating efforts to obtain stable, oligomer-enriched preparations [8, 9]. To address these challenges, nanoparticle-assisted platforms have been developed to stabilize amyloid oligomers at plasmonic nanoparticle interfaces for oligomer-targeted drug screening; however, these systems rely on nanoparticle templating and do not directly control the intrinsic aggregation pathway of proteins in solution [10–13]. For example, engineered nanoparticle interfaces can modulate amyloid conformational transitions through surface-mediated interactions, yet such approaches remain inherently dependent on interfacial templating effects rather than solution-phase pathway control [14]. Additionally, temperature- and time-controlled protocols have enabled partial enrichment of oligomeric species; however, such approaches rely on the intrinsic progression of aggregation and do not allow precise or reproducible control over specific intermediate states. As a result, they are largely limited to short peptide fragments and do not provide a mechanically programmable strategy for whole-protein amyloid formers [15, 16].

From a kinetic perspective, amyloid assembly proceeds through coupled microscopic steps—including primary nucleation (*k*_*n*_), fibril elongation (*k*_+_), and secondary nucleation (*k*_2_)—that collectively determine the time-dependent aggregate mass *M*_(*t*)_ [17–19]. In conventional systems, amplification through fibril elongation and secondary nucleation drives rapid fibril maturation, thereby limiting sustained oligomer accumulation. Although numerous thermal, mechanical, biological, and chemical strategies have been developed to regulate amyloid assembly [20–25], these approaches have largely focused on controlling fibril growth rather than on the deliberate and scalable production of oligomer-rich, fibril-suppressed assemblies.

Recent advances in amyloid kinetics have demonstrated that aggregation is not governed solely by solution chemistry but is also highly sensitive to mechanical and interfacial environments, giving rise to a mechanochemical perspective on amyloid assembly [21, 26–30]. In particular, air–water interfaces (AWI) can function as catalytic surfaces that promote primary nucleation and rebalance competing microscopic steps by locally increasing protein concentration and lowering the effective free-energy barrier for nucleation [21, 31, 32]. Hydrodynamic shear modulates molecular transport, interfacial residence time, fibril fragmentation probability, and aggregate detachment [21, 27, 33].

These effects collectively reweigh the reaction network, altering whether monomers preferentially proceed toward fibrillar elongation or accumulate in oligomeric states. Under shear flow, the local shear rate 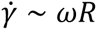 (where *ω* is angular velocity and *R* is radial position) determines the viscous stress 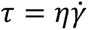, which can mechanically perturb elongating fibrils and alter effective kinetic rate constants [27, 34]. Rather than treating agitation and interfacial phenomena as experimental artifacts, these mechanochemical inputs can therefore be rationally engineered to bias kinetic partitioning in a bottom-up manner. By deliberately controlling the hydrodynamic flow fields and AWI exposure, researchers can bias early nucleation events while suppressing fibril growth, thereby shifting the assembly landscape toward oligomer-rich fibril-suppressed outcomes. Although shear forces and interfacial effects are known to modulate nucleation and growth kinetics, their deliberate application to bias assembly toward oligomer-dominant states has not been systematically achieved [27, 34, 35]. In most experimental settings, mechanical perturbations such as orbital shaking, stirring, or bead-assisted agitation are introduced primarily to shorten lag phases or enhance assay reproducibility [27, 35–39]. These approaches typically increase mass transport, promote fibril fragmentation into seeding-competent species, and amplify secondary nucleation on fibril surfaces—collectively accelerating fibrillization rather than suppressing it [18, 27]. Moreover, such perturbations are often implemented in poorly defined hydrodynamic regimes, where shear intensity, interfacial renewal, and centrifugal effects are neither quantified nor independently controlled. Consequently, mechanical inputs largely function as kinetic accelerants rather than as programmable variables for pathway selection. Therefore, a controllable, sustained mechanical platform capable of reproducibly promoting oligomer accumulation while limiting fibril propagation is required.

In the present study, we translated this mechanochemical framework into a practical and scalable strategy for oligomer-selective amyloid synthesis. We introduce a physical platform based on high-speed axial rotation implemented using a thermo-axial rotator to impose well-defined hydrodynamic and interfacial conditions without exogenous chemical additives. Continuous axial rotation generates sustained shear forces and centrifugal acceleration, thereby reshaping the aggregation landscape through a controlled mechanical input. Using hen egg-white lysozyme (HEWL) as a model amyloid-forming protein, we demonstrated that this regime selectively reroutes the assembly directly from monomeric precursors toward oligomer-rich states, while suppressing productive fibril formation and subsequent elongation.

To the best of our knowledge, this study provides the direct experimental evidence that sustained and quantitatively tunable mechanical control can bias an amyloid-forming system toward oligomer-enriched assembly states while effectively suppressing fibrillation. This chemical-free, bottom-up mechanochemical strategy establishes a reproducible route to oligomer-enriched assemblies, enabling the systematic investigation of structure–toxicity relationships and facilitating the development of oligomer-selective interventions, while also opening new opportunities for functional amyloid-based nanomaterial design [40].

## 2. RESULTS AND DISCUSSION

### 2.1. Hydrodynamic Regime Mapping and Mechanochemical Modulation of Amyloid Assembly

To establish a mechanically programmable platform for amyloid assembly, we first characterized the hydrodynamic environment generated by the axial rotation (Figure 1a). Under static conditions (0 RPM), the solution remained diffusion-limited, with mass transport governed primarily by Brownian diffusion and gravitational settling. No coherent rotational flow was generated, and intermolecular encounters occurred stochastically within the bulk phase.

**Figure 1.**
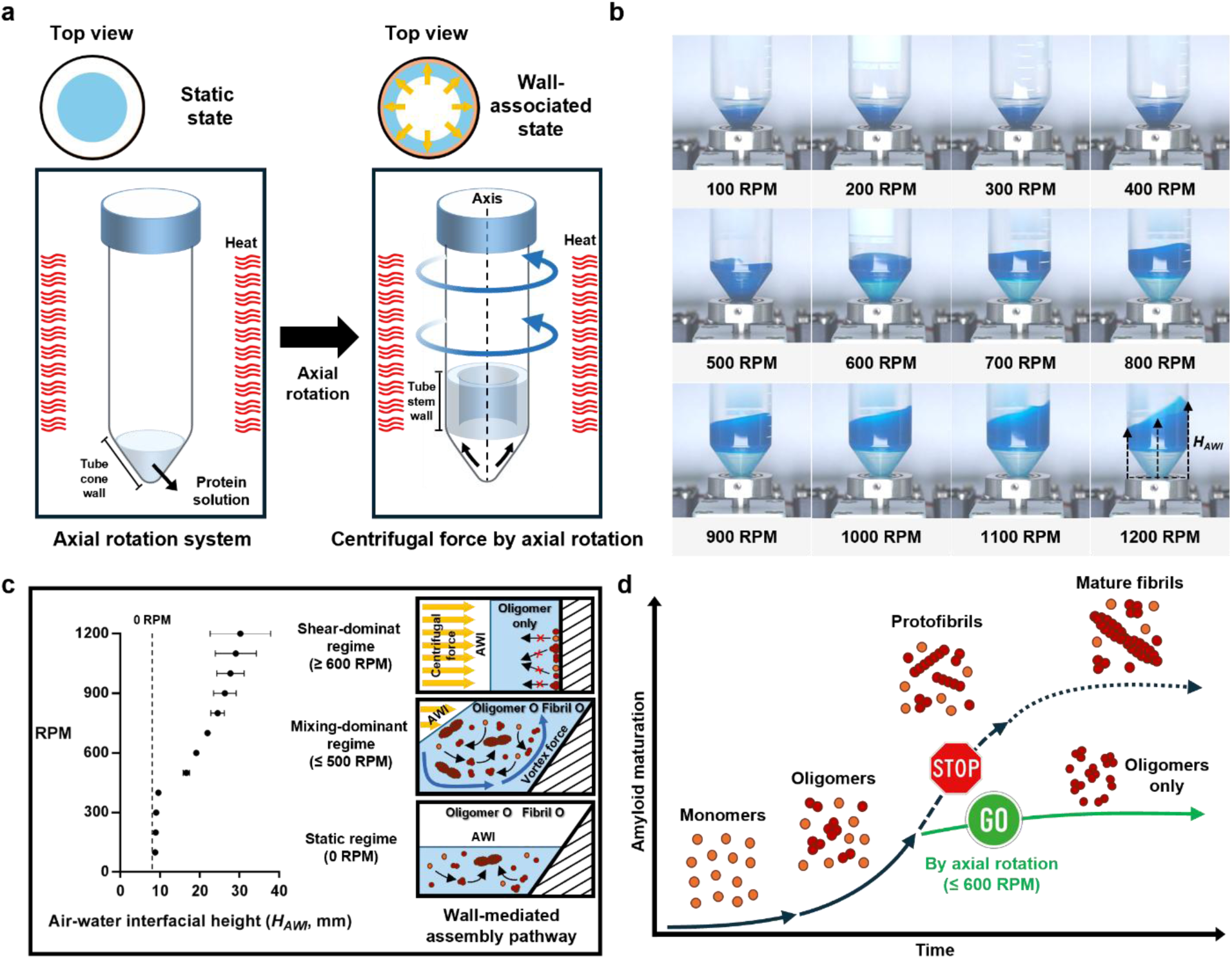
Concept and mechanochemical modulation of amyloid assembly via axial rotation. (a) Schematic illustration of the axial rotation setup. A conical tube was rotated along its longitudinal axis. Under static conditions (0 RPM), no coherent rotational flow is generated. Upon axial rotation, circumferential wall-associated flow develops, restructuring the internal hydrodynamic environment. (b) Representative images of the solution inside the tube at rotational speeds from 100 to 1200 RPM. Increasing RPM progressively tilts the air–water interface (AWI) along the tube wall due to centrifugal forces. (c) Quantification of the AWI height (H_AWI_, mm) as a function of rotational speed. H_AWI_ was defined as the maximum vertical displacement of the AWI measured along the inner wall of the tube. Data are presented as the mean ± SD (n = 3). Inset schematic illustrations depict distinct hydrodynamic regimes corresponding to low-RPM (diffusion-dominant), intermediate-RPM (mixing-dominant), and high-RPM (shear-dominant) conditions. (d) A conceptual model illustrating the proposed mechanochemical regulation of amyloid assembly pathways. Below a critical rotational threshold (≤600 RPM), aggregation follows a fibril-amplifying pathway. Above this threshold, fibril elongation is suppressed, and oligomer-dominant assemblies emerge.

Upon axial rotation, the hydrodynamic regime was fundamentally restructured. The conical tube rotates along its longitudinal axis, producing a circumferential flow field adjacent to the inner wall. Unlike orbital shaking, which induces a spatially heterogeneous flow field [41], axial rotation imposes a symmetry-constrained flow profile in which the centrifugal acceleration and wall-associated shear are directly tunable by the rotational speed (RPM). The centrifugal acceleration scales as *a*_*c*_ = *ω*^2^*R*, while the local shear rate near the wall scales approximately as 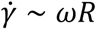, enabling quantitative modulation of viscous stress and interfacial deformation.

As the rotational speed increases, distinct interfacial deformations become evident (Figure 1b). The AWI progressively tilts along the inner wall owing to centrifugal forces, reflecting the redistribution of fluid mass and establishment of radial pressure gradients. Because the interfacial morphology depends on the sample volume, we evaluated the 1, 2, and 5 mL conditions and selected 1 mL as the reference volume, which exhibited the most pronounced and reproducible interfacial displacement (Figure S1).

To quantify this transition, we defined the AWI height (*H_AWI_*) as the maximum vertical displacement of the interface (Figure 1c). The *H_AWI_* increased progressively with RPM, confirming that axial rotation induced a controllable and reproducible modification of the internal flow field. The variability across replicates remained minimal at low-to-moderate RPM but increased at higher rotational speeds, likely reflecting the greater sensitivity of the interfacial configuration under strong centrifugal force.

Under static conditions (0 RPM) or low rotational speeds (<350 RPM), the centrifugal displacement was limited, and the resulting *H_AWI_* remained small. Consequently, protein species experience prolonged residence near the walls and AWI. Under fully static conditions, intermolecular encounters arise primarily from Brownian motion and diffusion-driven molecular mixing within the bulk solution, whereas gravitational settling further increases the local concentration near the solid boundaries. This diffusion-limited environment favors heterogeneous nucleation at interfaces and promotes subsequent fibril elongation, resulting in fibril-dominant assemblies [42]. In contrast, at higher rotational speeds (≥600 RPM), centrifugal forces strongly deform the AWI, resulting in an increased *H_AWI_* and elevated wall shear. These conditions shorten the interfacial residence time and mechanically perturb the elongating fibrils, thereby suppressing fibril extension and biasing the system toward oligomer-dominant assemblies.

As HEWL did not undergo amyloid formation at room temperature, aggregation was performed at 60 °C in sealed conical tubes. Importantly, the surface temperature remained stable across the RPM conditions (Figure S2), indicating that the RPM-dependent structural redistribution cannot be attributed to temperature differences but instead reflects the mechanical effects introduced by axial rotation.

The conceptual framework summarized in Figure 1d integrates these observations. Below a critical rotational threshold, aggregation proceeded along a fibril-amplifying pathway dominated by elongation and surface-mediated nucleation. Above this threshold, mechanically imposed shear and centrifugal restructuring suppress extended fibril formation and bias the system toward oligomer stabilization. Therefore, axial rotation functions not only as a mixing modality but also as a quantitatively tunable mechanochemical regulator of amyloid pathway selection. This hydrodynamic regime mapping established the physical foundation for the structural and kinetic analyses presented below.

### 2.2. RPM-Dependent Redistribution of Amyloid Assembly States

To determine whether axial rotation redefines the structural outcome of amyloid assembly, we quantified β-sheet formation and hydrophobic surface exposure using thioflavin T (ThT) and 8-anilinonaphthalene-1-sulfonic acid (ANS), respectively (Figure 2a) [43]. ThT fluorescence, indicative of β-sheet–rich fibrillar structures, increased from static conditions to 350 RPM, suggesting enhanced fibril formation under moderate rotational input. Beyond 350 RPM, however, the ThT intensity declined sharply and remained suppressed at higher speeds. In contrast, ANS fluorescence—reporting solvent-exposed hydrophobic surfaces, typically associated with partially folded or oligomeric intermediates, remained comparatively stable or were modestly elevated at high RPM. This opposing trend indicated a redistribution of structural populations rather than a reduction in total aggregation. Consistent with this interpretation, Bicinchoninic Acid (BCA) analysis confirmed no statistically significant differences in the total protein concentration across the RPM conditions (Figure S3), demonstrating that fluorescence changes reflect structural reweighting rather than differences in aggregate mass.

**Figure 2.**
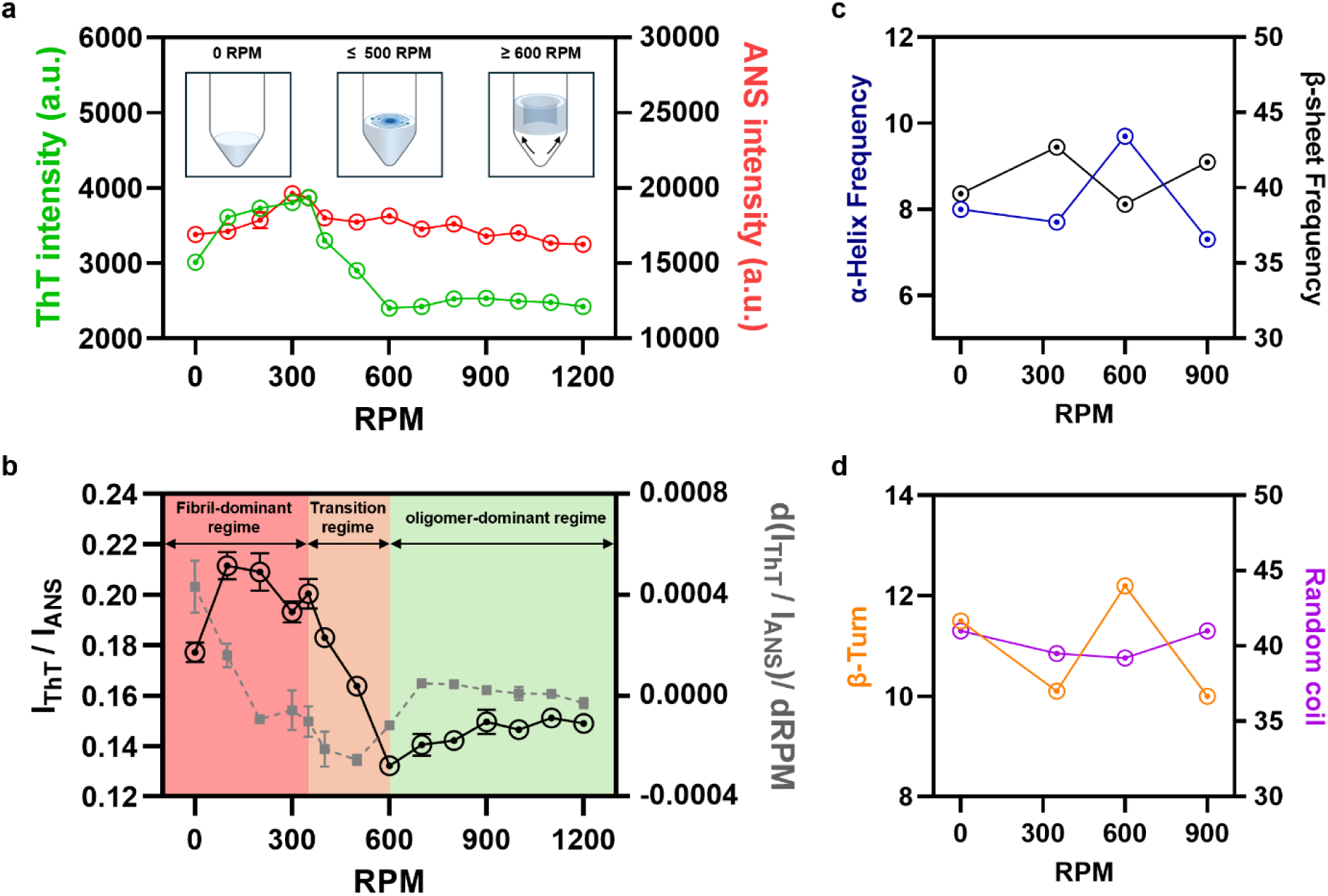
RPM-dependent modulation of amyloid assembly states. (a) Fluorescence responses of thioflavin T (ThT) and 8-anilinonaphthalene-1-sulfonic acid (ANS) as a function of rotational speed. ThT intensity (green, left axis) and ANS intensity (red, right axis) were measured under identical experimental conditions and plotted against RPM. Representative schematics illustrate the hydrodynamic states of the solution under static, intermediate, and high-RPM conditions. Data are the mean ± SD (n ≥ 3). (b) RPM-dependent variation of the ThT/ANS fluorescence intensity ratio. The ratio serves as an indicator of the relative contribution of fibrillar versus oligomeric assemblies. Regime classification was performed by first deriving the normalized ThT/ANS ratio with respect to RPM, identifying three regimes: a fibril-dominant regime (0–350 RPM), a high-slope transition regime (350–600 RPM), and an oligomer-dominant regime (>600 RPM). (c, d) Secondary structure analysis derived from circular dichroism (CD) spectroscopy, showing α-helix, β-sheet, β-turn, and random coil content under representative RPM conditions.

To quantify the relative balance between β-structured assemblies and oligomeric species, the ThT/ANS fluorescence ratio was used as a composite metric (Figure 2b). Higher ratios indicate β-structure-dominant assemblies, whereas lower ratios reflect oligomer-dominant assemblies. Based on the first derivative of the ThT/ANS ratio as a function of rotational speed (Figure 2b), three assembly regimes were defined:

(i) low-RPM plateau regime (0–350 RPM), characterized by high ThT/ANS ratios and pronounced β-sheet formation; (ii) high-slope transition regime (350–600 RPM), marked by rapid decline in the ratio and coexistence of fibrillar and oligomeric populations; and (iii) high-RPM plateau regime (>600 RPM), in which the ratio remained suppressed, indicating enrichment of oligomeric assemblies relative to mature fibrils.

Importantly, this regime classification reflects structural balance rather than aggregation efficiency. Therefore, axial rotation does not simply attenuate fibril formation; instead, it redefines the relative populations of structurally distinct amyloid species in a speed-dependent manner.

Circular dichroism (CD) spectroscopy independently confirms this structural redistribution (Figures 2c and 2d and S4). Low-RPM samples exhibited elevated β-sheet content consistent with fibril-dominant assemblies (Figure 2c). At 600 RPM, β-sheet fraction was markedly reduced and accompanied by an increased β-turn contribution, indicating enrichment of partially ordered conformations consistent with oligomeric assemblies (Figure 2d) [44].

Notably, 900 RPM samples showed partial recovery of β-sheet signal relative to 600 RPM despite residing within the oligomer-dominant fluorescence regime. This observation suggests that high-RPM conditions do not maintain a structurally disordered state but instead promote the formation of compact oligomeric assemblies containing β-structural elements distinct from those of mature fibrils.

Collectively, these data demonstrate that axial rotation induces a discrete structural transition within the 350–600 RPM regime, beyond which fibrillar structures are reduced, and oligomer-dominant conformational ensembles emerge. Moreover, the 600 and 900 RPM assemblies were not structurally identical but corresponded to distinct oligomeric assemblies formed under different mechanochemical conditions.

### 2.3. Morphological Validation of RPM-Dependent Assembly States by Atomic Force Microscopy

To directly validate the structural redistribution inferred from the fluorescence and CD analyses, we performed atomic force microscopy (AFM) imaging at representative rotational speeds (Figure 3). This analysis enabled discrimination between the elongated fibrillar structures and discrete oligomeric particles and provided quantitative height distributions to complement the spectroscopic data. Under static conditions (0 RPM), abundant elongated fibrils were observed, exhibiting micrometer-scale lengths and a well-defined linear morphology (Figure 3a). Height profiles extracted along representative fibrils showed distinct nanometer-scale peaks consistent with mature amyloid structures. At 200 and 350 RPM, the fibrillar density increased, and the fibrils appeared more interconnected, suggesting enhanced fibril growth under moderate rotational input (Figure 3b).

**Figure 3.**
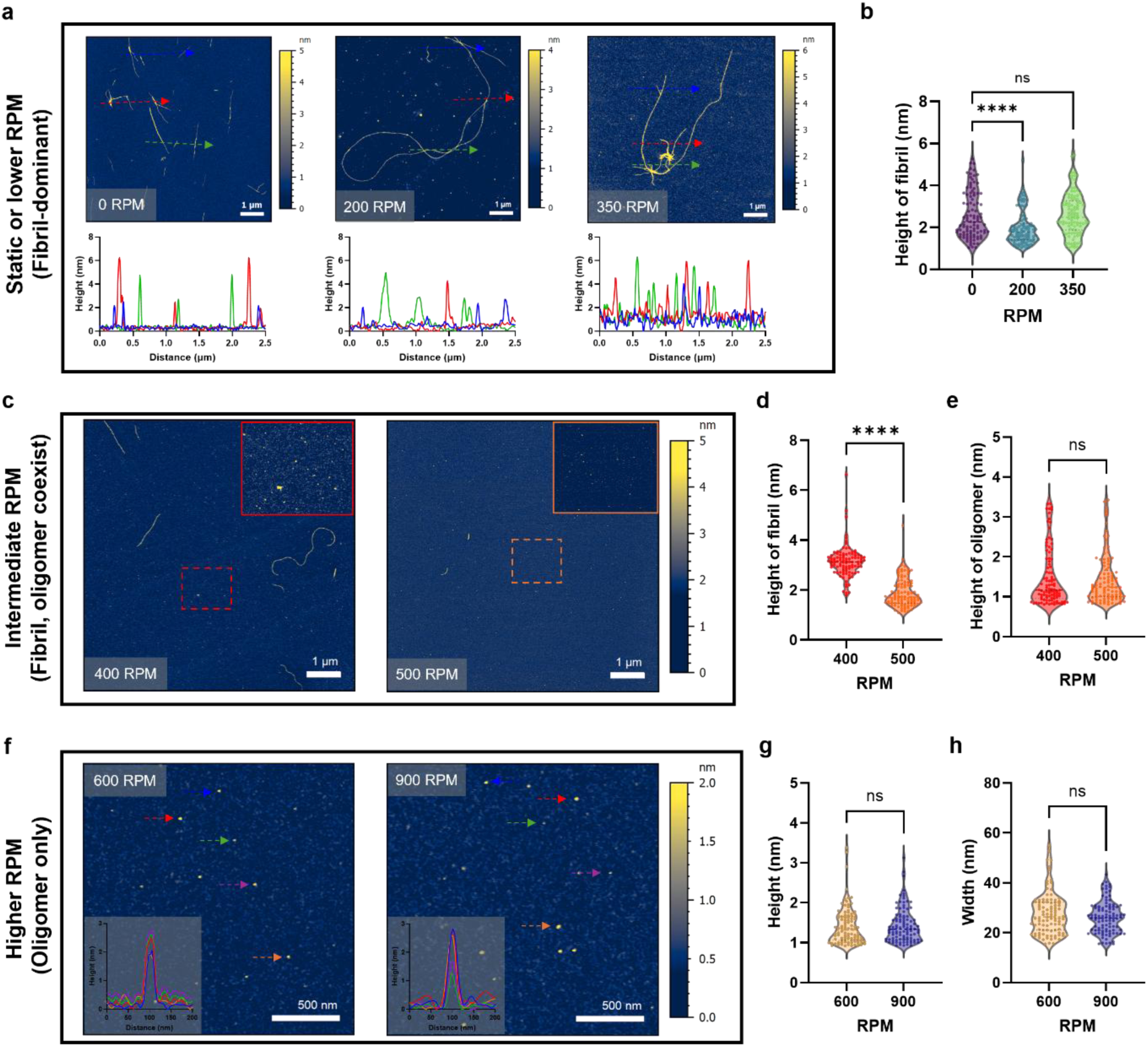
AFM analysis validating RPM-dependent morphological transitions of amyloid assemblies formed at 60 °C for 3 days. (a) Representative AFM height images of samples prepared at 0, 200, and 350 RPM, corresponding to the fibril-dominant regime. Cross-sectional profiles extracted along colored arrows are shown below each image. Scale bars: 1 µm. (b) Violin plots summarizing fibril height distributions measured from AFM images in (a), confirming increased fibrillar dimensions under low-to-moderate RPM conditions. (c) AFM images of samples prepared at 400 and 500 RPM, corresponding to the transition regime. Magnified insets (dashed boxes) highlight representative oligomer-rich regions coexisting with attenuated fibrils. Scale bars: 1 µm. (d, e) Violin plots summarizing height distributions of fibrils and oligomers under intermediate RPM conditions (400 and 500 RPM), illustrating suppression of fibril elongation without significant change in oligomer size. (f) High-magnification AFM images of assemblies prepared at 600 and 900 RPM, corresponding to the oligomer-dominant regime. Elongated fibrils are largely absent, and uniformly distributed spherical particles predominate. Insets show representative height profiles. Scale bars: 500 nm. (g, h) Violin plots summarizing oligomer height and lateral width distributions under high-RPM conditions (600 and 900 RPM). No statistically significant differences were observed between these conditions. Data represent measurements of individual particles (n = 100 per condition) obtained from multiple independent images.

A distinct morphological transition emerged at intermediate rotational speeds (400–500 RPM) (Figure 3c). The fibrils became progressively shorter and thinner, indicating the suppression of longitudinal elongation and lateral thickening. Simultaneously, numerous discrete nanoscale particles were detected. Height analysis revealed that the oligomeric particle size remained statistically comparable between 400 and 500 RPM (Figure 3d,e), suggesting that this transition regime primarily reflects the attenuation of fibril architecture rather than structural diversification of the oligomer population.

At higher rotational speeds (600 and 900 RPM), elongated fibrils were largely absent and were replaced by uniformly distributed spherical assemblies (Figure 3f and S5). High-resolution AFM scans revealed narrow height distributions consistent with oligomeric assemblies. Quantitative analysis confirmed that the particle height and lateral width showed no statistically significant differences between the 600 and 900 RPM samples (Figure 3g,h).

These morphological findings were consistent with the fluorescence-defined assembly regimes (Figure 2). Under low RPM conditions, fibril-dominant assemblies were observed, whereas intermediate rotational speeds produced mixed populations containing both fibrillar and particulate structures. At about 600 RPM, the system shifts toward oligomer-dominant states characterized by dispersed nanoscale particles. Importantly, although the 600 and 900 RPM samples exhibited similar nanoscale morphologies, CD spectroscopy revealed distinct secondary structural signatures (Figure 2c,d). In particular, the 600 RPM assemblies displayed reduced β-sheet content accompanied by increased β-turn contributions, whereas the 900 RPM samples showed partial recovery of β-sheet structure. These spectroscopic differences indicate that the oligomeric assemblies generated at 600 and 900 RPM represent mechanochemically differentiated states despite their comparable particle dimensions [45, 46].

Taken together with fluorescence and spectroscopic analyses, AFM observations provide morphological validation of the RPM-dependent structural transition identified in Figure 2. Specifically, the fibrillar structures diminish across the transition regime and are replaced by dispersed spherical assemblies at higher rotational speeds. These results demonstrate that RPM-dependent mechanical input redefines the amyloid assembly pathway and generates distinct aggregate morphologies across rotational regimes.

### 2.4. Seeding Effects of RPM-Defined Assemblies on Amyloid Aggregation Kinetics

Given that an oligomer-like morphology does not necessarily imply functional seeding competence, we examined whether the assemblies generated by axial rotation acted as effective seeds during amyloid aggregation under conventional heating conditions. The preformed assemblies obtained under representative rotational conditions were mixed in a 1:1 ratio with fresh HEWL monomers, and ThT fluorescence was monitored under identical incubation conditions (Figure 4a–c).

**Figure 4.**
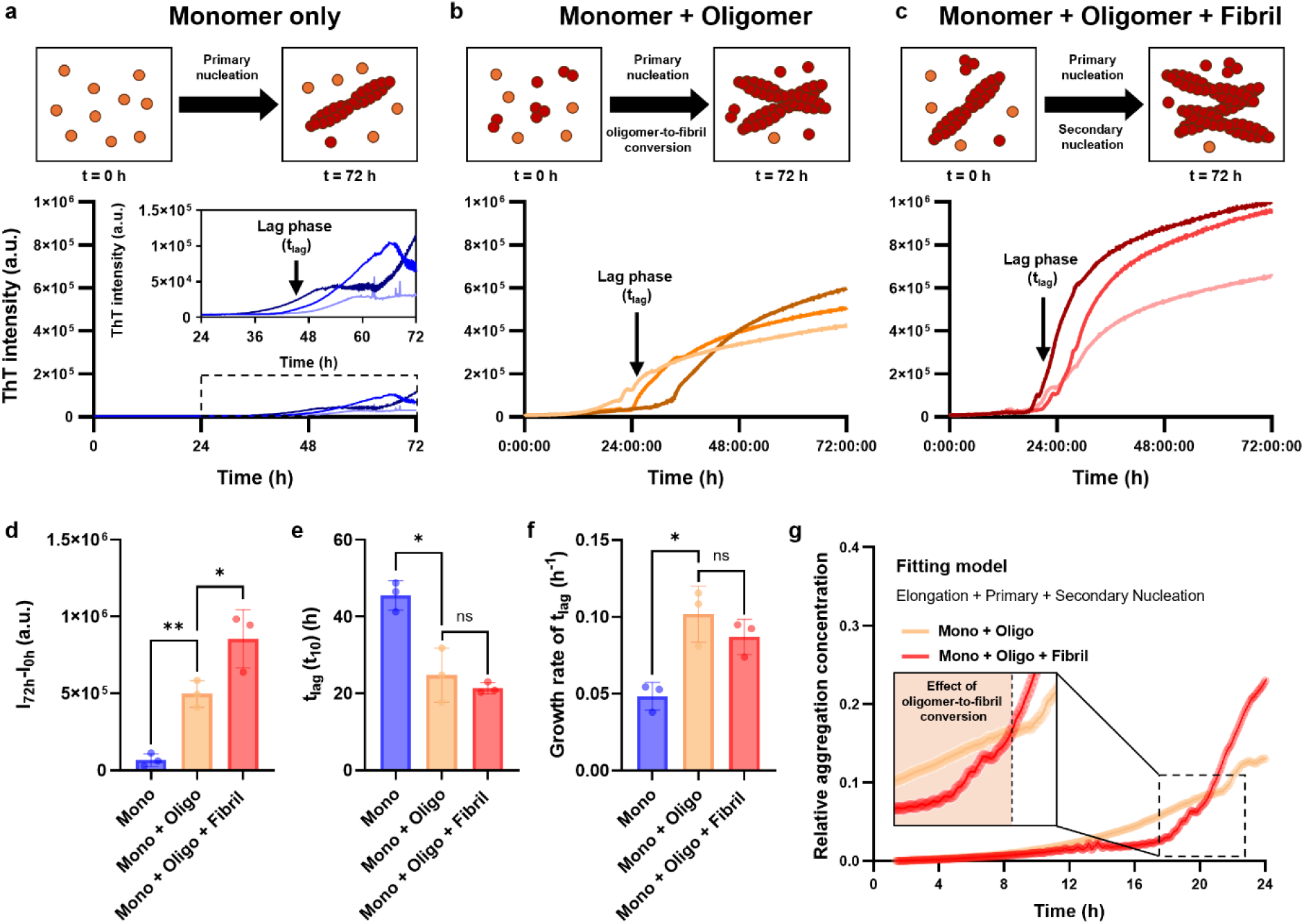
Seeding effects of RPM-defined assemblies on amyloid aggregation kinetics. (a–c) Schematic illustration and corresponding ThT fluorescence kinetics of (a) the HEWL monomer alone, (b) the monomer mixed 1:1 with oligomer-dominant assemblies prepared at 600 RPM, and (c) the monomer mixed 1:1 with assemblies prepared at 350 RPM containing both oligomers and fibrils. The aggregation was monitored under identical conditions. The curves represent independent replicates. (d–f) Quantitative parameters extracted from the kinetic traces in (a–c), including the (d) maximum fluorescence intensity. (e) Early aggregation onset quantified using t₁₀, defined as the time required to reach 10% of the maximum fluorescence intensity [50]. (f) Apparent growth rate at the lag-to-growth transition (mean ± SD); the 600 RPM condition shows a higher mean value but is not statistically significant. (g) Global kinetic fitting was performed using the AmyloFit platform, based on a model that incorporates elongation, primary nucleation, and secondary nucleation. The enlargement of the early time window highlights the differences in the initial aggregation behavior between oligomer- and fibril-containing assemblies. Data are presented as the mean ± SD from independent experiments (n = 3). Statistical significance was evaluated using one-way ANOVA followed by Tukey’s multiple comparison test (p < 0.05).

Amyloid aggregation generally follows nucleation-dependent polymerization kinetics, in which primary nucleation generates initial aggregates that subsequently grow through elongation and secondary nucleation [18, 47–50]. Consistent with this framework, the monomer alone exhibited a delayed onset of ThT fluorescence, followed by a gradual increase in signal (Figure 4a). When mixed with oligomer-dominant assemblies generated at 600 RPM (Figure 4b), the lag phase was modestly reduced and the fluorescence signal increased earlier than in the monomer condition, indicating accelerated early stage aggregation kinetics induced by oligomeric seeds. This kinetic acceleration became more pronounced when the assemblies generated at 350 RPM, containing both fibrils and oligomers, were introduced (Figure 4c). In this case, the lag phase was significantly shortened, the growth phase steepened, and the maximum fluorescence intensity increased substantially. These features indicate enhanced fibril templating and amplification, consistent with secondary nucleation on preexisting fibrillar surfaces [49, 51, 52].

Quantitative parameter extraction (Figure 4d–f) further delineated these differences. The maximum fluorescence intensity was the highest for the fibril-containing (350 RPM) condition (Figure 4d), reflecting increased total fibril accumulation. In contrast, analysis of the time required to reach a 10-fold increase in initial fluorescence (*t₁₀*) revealed stronger early-stage acceleration in the oligomer-dominant (600 RPM) condition (Figure 4e).

The apparent growth rate at the lag-to-growth transition showed a higher mean value for the 600 RPM-derived assemblies compared to the 350 RPM condition, although this difference was not statistically significant (Figure 4f). Under conditions in which the total protein concentration was identical, this directional trend may reflect a greater contribution of oligomeric species to early aggregation events in the 600 RPM sample. To further examine these kinetic differences, global fitting was performed using the AmyloFit platform with a combined elongation, primary nucleation, and secondary nucleation model (Figure 4g) [19]. Enlargement of the early time window revealed a steeper initial rise for oligomer-containing samples, whereas fibril-containing assemblies exhibited stronger signal amplification during later stages of aggregation. Collectively, these observations suggest that oligomer-rich assemblies tend to influence the early aggregation kinetics, whereas fibril-containing assemblies promote amplification more effectively during subsequent growth.

Importantly, although early kinetic trajectories differed between the oligomer-rich assemblies and fibril–oligomer mixtures, AFM analysis of the 72-h end-point samples (Figure S6) revealed elongated fibrils in both cases. Consistent with this observation, AFM imaging also detected a small number of relatively short fibrils in samples prepared under static conditions despite identical sampling concentrations. These results suggest that the distinct aggregation kinetics arise primarily from differences in the aggregate composition of the initial seeds rather than from differences in their ability to form fibrillar structures during prolonged incubation. Collectively, these results demonstrate that assemblies generated by axial rotation not only retain seeding capability, but also impose distinct kinetic biases on subsequent aggregation, depending on their oligomer–fibril composition.

### 2.5. Cytotoxic Profiles of RPM-Defined Assemblies in SH-SY5Y Cells

To determine whether the structurally distinct HEWL assemblies generated under different rotational regimes elicited different biological responses, we evaluated their cytotoxicity in SH-SY5Y neuroblastoma cells (Figure 5). HEWL samples incubated at 60 °C for 3 days under 0, 350, 600, and 900 RPM were applied at 1 and 7 μM, and cellular responses were assessed morphologically and biochemically.

**Figure 5.**
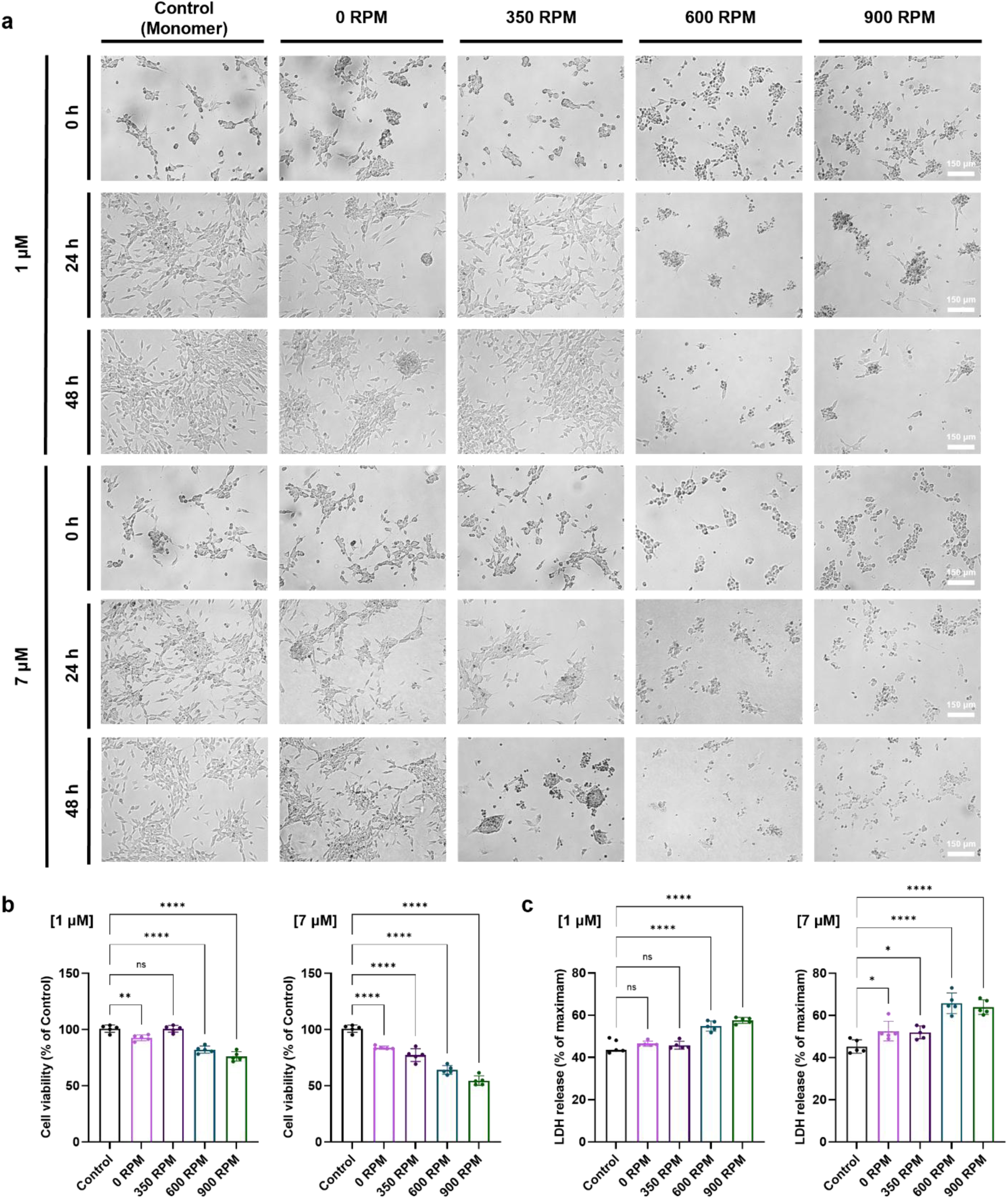
Cytotoxic effects of RPM-defined HEWL assemblies on SH-SY5Y cells. HEWL samples (monomer control and assemblies generated at 0, 350, 600, and 900 RPM) were applied to SH-SY5Y cells at concentrations of 1 and 7 μM. (a) Representative bright-field images of SH-SY5Y cells after treatment with monomers and RPM-defined assemblies. Images were acquired at defined post-treatment time points. Scale bars: 150 μm. (b) Cell viability was assessed using the Cell Counting Kit-8 assay. Data are presented as the mean ± SD (n = 5). (c) Cytotoxicity was evaluated using a lactate dehydrogenase (LDH) release assay under the same conditions. Data are presented as the mean ± SD (n = 5). Statistical comparisons were performed relative to the untreated control cells.

At 1 μM, bright-field imaging revealed modest condition-dependent differences in cell morphology (Figure 5a). Cells treated with the monomer or low-RPM assemblies (0 and 350 RPM) maintained their adherent morphology and confluency. In contrast, 600 and 900 RPM samples induced visible cell clustering and partial detachment, suggesting altered cell–cell or cell–substrate interactions under oligomer-dominant conditions [53, 54]. At 7 μM, morphological disruption became pronounced across all aggregate-treated groups, with reduced confluency and increased cell rounding. These findings indicate concentration-dependent amplification of aggregate-induced cellular stress.

Quantitative viability testing using the cell counting kit-8 (CCK-8) assay confirmed these trends (Figure 5b). At 1 μM, all aggregate-treated groups exhibited reduced metabolic activity relative to monomer-treated controls, with the greatest decrease observed for assemblies generated at 600 and 900 RPM. At 7 μM, viability was substantially reduced across conditions, but oligomer-dominant samples (≥600 RPM) maintained comparatively lower metabolic activity than fibril-containing assemblies.

Measurement of lactate dehydrogenase (LDH) release further clarified the effects on membrane integrity (Figure 5c). At 1 μM, membrane damage was detectable but moderate. At 7 μM, LDH release increased significantly, particularly for low-RPM (0 and 350 RPM) samples containing fibrillar structures. Notably, despite pronounced clustering and detachment induced by the 600 and 900 RPM assemblies, LDH release did not increase proportionally.

The divergence between metabolic suppression and LDH release suggests a mechanistically distinct cytotoxic profile. Fibril-containing assemblies appeared to promote membrane disruption, which is consistent with lytic damage, whereas oligomer-dominant assemblies preferentially impaired cellular adhesion and metabolic function before overt membrane rupture.

Taken together, these results demonstrated that mechanochemically defined structural states translate into distinct cytotoxic phenotypes. Assemblies enriched in oligomeric species (≥600 RPM) exhibit enhanced metabolic toxicity and morphological perturbation, whereas fibril-containing assemblies exert stronger membrane-disruptive effects at higher concentrations. Thus, axial rotation not only redefines the structural and kinetic properties of amyloid assemblies but also modulates their downstream cellular impact in a regime-dependent manner.

### 2.6. Live/Dead Response and Cytoskeletal Remodeling Induced by RPM-Defined Assemblies

To further examine the structural consequences of the RPM-defined assemblies on cellular integrity, we performed live/dead staining and cytoskeletal analysis of SH-SY5Y cells following exposure to samples generated under different rotational conditions (Figure 6).

**Figure 6.**
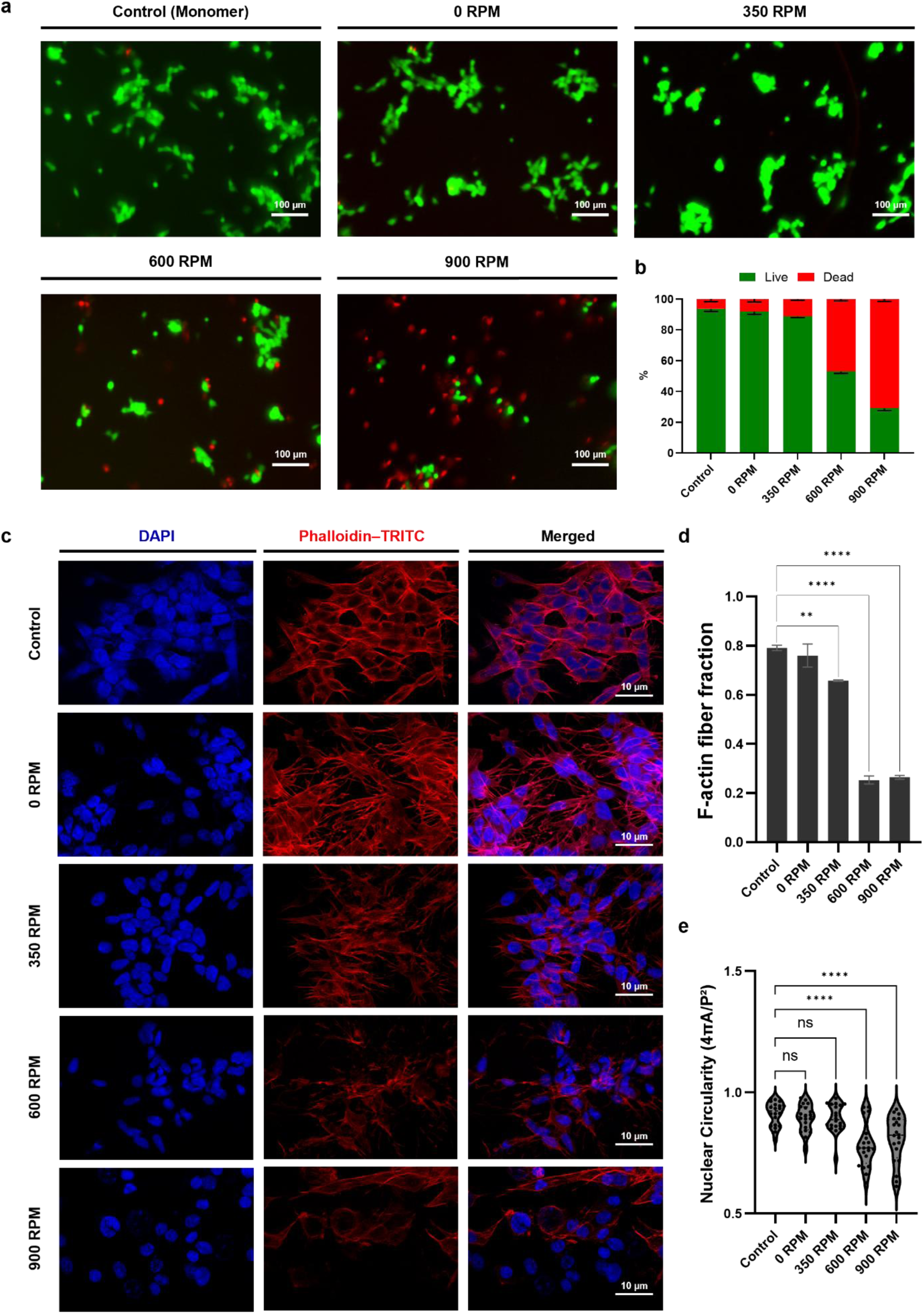
Live/dead response and cytoskeletal remodeling in SH-SY5Y cells treated with RPM-defined HEWL assemblies. HEWL samples (monomer control and assemblies prepared at 0, 350, 600, and 900 RPM) were applied to SH-SY5Y cells at a concentration of 1 μM. (a) Representative fluorescence images from live/dead staining after 1 h treatment with HEWL samples. Live cells are shown in green and dead cells in red. Scale bars: 100 μm. (b) Quantification of live and dead cell populations expressed as percentages of total cells. (c) Immunofluorescence staining of nuclei and F-actin cytoskeleton following 1 h treatment of HEWL samples and immediate fixation. Nuclei were stained with 4′,6-diamidino-2-phenylindole (blue), and F-actin was stained with phalloidin–TRITC (red). Merged images are shown. Scale bars: 10 μm. (d) Quantitative analysis of F-actin organization based on fluorescence intensity and F-actin–positive area fraction. (e) Distribution of nuclear circularity metrics presented as violin plots.

At 7 μM, cell death exceeded 96% across aggregate-treated conditions (Figure S7), precluding meaningful structural analysis. Therefore, cytoskeletal remodeling was evaluated at 1 μM, where differential phenotypes were observable. Live/dead staining at 1 μM revealed regime-dependent viability differences (Figure 6a,b). The monomer-treated cells predominantly exhibited green fluorescence, indicating high cell viability. Assemblies generated at 0 and 350 RPM induced moderate increases in red fluorescence. In contrast, the 600 and 900 RPM samples showed a marked increase in the dead cell population relative to the low-RPM conditions, consistent with the enhanced metabolic suppression observed in Figure 5. Notably, the magnitude of cell death observed in the live/dead assay was greater than that inferred from the CCK-8 and LDH measurements. This difference likely reflects the distinct assay readouts, as the live/dead assay directly reports membrane integrity, whereas the CCK-8 and LDH assays primarily capture metabolic activity and bulk enzyme release.

Immunofluorescence staining of F-actin and nuclei provided structural insights into the cellular perturbations associated with HEWL assembly (Figure 6c). The monomer-treated cells displayed a well-spread morphology with dense, continuous F-actin networks and organized cortical structures. Cells treated with 350 RPM assemblies showed partial disruption of filament organization, but largely preserved cytoskeletal continuity. In contrast, the 600 and 900 RPM assemblies induced pronounced cytoskeletal destabilization, which was characterized by reduced F-actin density, fragmented filament networks, and impaired cell spreading. The cells exhibited rounded morphology with diminished actin-rich protrusions, indicating compromised structural integrity.

The quantitative analysis confirmed these observations (Figure 6d). F-actin–positive area fraction and fluorescence intensity were significantly reduced under ≥600 RPM conditions compared to monomer and 350 RPM treatments. This reduction reflected the substantial attenuation of filamentous actin organization under oligomer-dominant conditions.

The nuclear morphology analysis further supported this distinction (Figure 6e). While low-rpm treatments produced relatively narrow circularity distributions, the 600 and 900 RPM samples exhibited broadened distributions and increased deviations from the baseline nuclear shape metrics. These alterations are consistent with cytoskeletal collapse–associated mechanical stress and impaired structural coupling between the actin cytoskeleton and the nucleus.

Taken together, these data demonstrate that oligomer-enriched assemblies generated at high rotational speeds are associated with more pronounced intracellular structural disruption than fibril-containing assemblies. Importantly, this cytoskeletal remodeling occurs at submaximal cytotoxic concentrations (1 μM) and results in distinct cellular phenotypes.

## 3. CONCLUSION

In this study, we demonstrated that sustained axial rotation enables programmable mechanochemical control of amyloid assembly pathways. By imposing quantitatively tunable centrifugal acceleration and wall-associated shear using a thermal axial rotator, we systematically restructured the hydrodynamic environment without chemical additives, thereby reweighting the microscopic steps governing aggregation.

Hydrodynamic regime mapping revealed a discrete transition at approximately 600 RPM. Below this threshold, aggregation proceeded along a fibril-amplifying pathway characterized by enhanced β-sheet formation, elongated fibrillar morphology, and strong secondary nucleation–mediated amplification. Above this threshold (≥600 RPM), fibrillar growth was strongly attenuated, and oligomer-dominant assemblies predominated. Importantly, this transition did not reflect a reduction in aggregate mass but rather a redistribution of structural populations, as confirmed by fluorescence normalization, circular dichroism, and AFM analysis.

Functionally, these mechanochemically defined assemblies exhibited distinct kinetic behaviors. Fibril-containing species preferentially enhance secondary nucleation and aggregate amplification, whereas oligomer-dominant assemblies accelerate early stage aggregation without proportionally increasing the final fibril mass. Despite convergence toward comparable fibrillar architectures over extended incubation times, early kinetic partitioning was clearly modulated by rotational input.

This structural redistribution translates into a differential cellular phenotype. Oligomer-enriched assemblies generated at approximately 600 RPM induced greater metabolic suppression and pronounced cytoskeletal destabilization in SH-SY5Y cells, whereas fibril-containing assemblies exerted stronger membrane-disruptive effects at elevated concentrations. These findings establish a mechanochemical link among hydrodynamic boundary conditions, aggregation-pathway selection, and downstream cellular responses.

Collectively, our results suggest that axial rotation is a scalable, chemical-free platform for mechanochemical programming of amyloid assemblies. This approach provides a reproducible route to oligomer-enriched preparations and offers a controllable framework for probing structure–kinetic–toxicity relationships in amyloid-forming systems. While RPM-dependent modulation of assembly pathways was established, further studies are required to resolve the structural heterogeneity of oligomer populations generated under different rotational conditions and to elucidate the underlying mechanisms of their cytotoxicity. These aspects will be addressed in future work.

## 4. Experimental Section

### 4.1. Materials and Reagents

Hen egg-white lysozyme (HEWL; 10837059001), thioflavin T (ThT), and 8-anilinonaphthalene-1-sulfonic acid (ANS) were purchased from Sigma-Aldrich (USA) and were used without further purification. Phosphate-buffered saline (PBS) and hydrochloric acid were used for buffer preparation. SH-SY5Y human neuroblastoma cells were obtained from the Korea Cell Line Bank (Republic of Korea). Cell Counting Kit-8 (CCK-8) and CytoTox 96 Non-Radioactive Cytotoxicity Assay were used according to the manufacturer’s instructions. Unless otherwise stated, all other reagents were of analytical grade.

### 4.2. HEWL Monomer Preparation

HEWL powder was dissolved in deionized water to obtain a 2% (w/v; 20 mg mL⁻¹) solution, and the pH was adjusted to 2.0 using hydrochloric acid. The solution was filtered through a 0.20 μm polyvinylidene fluoride syringe filter (Whatman GD/X 25, Cytiva, USA) to remove pre-existing aggregates and was used as a monomeric HEWL stock solution. Aliquots (1 mL) were transferred to 50 mL conical tubes (Hyundai Micro Co., Ltd., Republic of Korea), tightly sealed with polytetrafluoroethylene tape to minimize evaporation, and mounted on a custom-built thermal axial rotator. Samples were incubated at 60 °C for 72 h under continuous axial rotation at designated speeds (0–1200 RPM). A 0 RPM sample incubated at 60 °C without rotation served as a static control, and the filtered HEWL stock was used as the reference representing the initial monomeric state.

### 4.3. ThT Fluorescence Assay

A ThT stock solution (10 mM) was prepared in water (pH 2.0) and diluted to a working solution (1 mM). HEWL assemblies were mixed with ThT to a final dye concentration of 10 μM and transferred to black 96-well plates [55]. Fluorescence was recorded using a multimode microplate reader (Synergy H1, BioTek, USA) equipped with Gen5 software in top-read mode (Ex 445 nm/Em 485 nm).

### 4.4. ANS Fluorescence Assay

ANS was added to samples to a final concentration of 20 μM and equilibrated for 5 min in the dark. The emission spectra (470–500 nm) were collected upon excitation at 350–380 nm (typical settings: Ex 370 nm/Em 480 nm) [43]. Increased ANS was interpreted as increased exposure of the hydrophobic surfaces within the aggregates.

### 4.5. Fluorescence Ratio Analysis

To evaluate the relative balance between the fibrillar and oligomeric assemblies, the ratio of ThT to ANS fluorescence intensity (ThT/ANS) was calculated at each RPM. The mean fluorescence values obtained from independent measurements were used for analysis. To identify RPM-dependent transitions in the assembly behavior, the first derivative of the ThT/ANS ratio with respect to the rotational speed was calculated using numerical differentiation. The derivative was estimated using a finite-difference approximation between adjacent RPM values (*dR*/*d*RPM ≈ (*R*_*i*+1_ − *R*_*i*_)/(RPM_*i*+1_ − RPM_*i*_)), where *R*represents the normalized ThT/ANS fluorescence ratio. Numerical differentiation was performed using GraphPad Prism software (GraphPad Software, USA).

### 4.6. Bicinchoninic Acid (BCA) Assay

The BCA assay was employed to determine the total protein concentration in the aggregated samples. Samples were diluted in deionized water (DW, pH 2.0) prior to analysis. The assay was performed according to the manufacturer’s protocol (23225, Thermo Fisher Scientific, USA), using bovine serum albumin as the standard for calibration. Absorbance was measured at 562 nm using a Synergy H1 multi-mode microplate reader (BioTek, USA), and protein concentrations were calculated based on the standard calibration curve.

### 4.7. CD Spectroscopy

To analyze secondary structure, CD spectra were obtained using a J-1500 spectropolarimeter (Jasco Corporation, Japan) at the Korea Basic Science Institute (KBSI, Ochang, Republic of Korea). For far-UV CD measurements, samples were diluted to 0.3 mg mL⁻¹ and measured in a 1 mm path-length quartz cuvette at 25 ℃ over the wavelength range of 190–260 nm. The corresponding buffer spectra were subtracted from the protein spectra, with each spectrum representing the average of multiple scans. Secondary structure contents (α-helix, β-sheet, β-turn, and random coil) were estimated using the Protein Secondary Structure Estimation module in Spectra Manager Version 2 software (Jasco Corporation).

### 4.8. ThT Aggregation Kinetics Assay

The amyloid aggregation kinetics were monitored using ThT fluorescence. HEWL assemblies generated under different rotational conditions were mixed with freshly prepared HEWL monomers at a 1:1 volume ratio. The final protein concentration was adjusted to the desired experimental condition, and ThT was added to a final concentration of 10 μM. Samples were transferred to black 96-well plates and incubated at 60 ℃. Fluorescence measurements were performed using a multimode microplate reader (Synergy H1, BioTek) in the kinetic mode with excitation at 445 nm and emission at 485 nm. Fluorescence intensity was recorded every 5 min for 72 h with orbital shaking.

To further analyze the aggregation kinetics, the ThT fluorescence traces obtained from the assay were directly used for kinetic modeling using the AmyloFit platform. The raw fluorescence intensity data were processed using the built-in preprocessing routine, where the baseline was corrected by averaging the initial three data points, and the curves were normalized to a plateau value of 1.0. The data were fitted using a model incorporating primary nucleation, elongation, and secondary nucleation without inhibition. Global fitting was performed across different conditions with selected parameters treated as shared or condition-specific variables, while certain parameters were constrained to avoid overparameterization and ensure stable convergence. The fitting analysis was used to compare relative kinetic trends among different assembly conditions rather than to assign uniquely identifiable microscopic rate constants.

### 4.9. AFM imaging

To visualize aggregate morphology, 20 μL of each sample was deposited onto freshly cleaved mica and incubated for 10 min at room temperature. The surface was gently rinsed twice with deionized water to remove unbound species and then dried under a gentle nitrogen stream. Atomic force microscopy (AFM) was performed in tapping mode using a MultiMode 8 atomic force microscope (Bruker, USA). Images were acquired using silicon cantilevers (Probes, PR-T300) under ambient conditions.

### 4.10. Cell Culture

Human neuroblastoma SH-SY5Y cells were maintained in high-glucose Dulbecco’s Modified Eagle’s medium (DMEM; Gibco, Thermo Fisher Scientific, USA) supplemented with 10% (v/v) fetal bovine serum and 1% penicillin–streptomycin (Thermo Fisher Scientific). The cells were cultured at 37 °C in a humidified incubator with 5% CO₂.

### 4.11. Cell Treatment

For cell-based assays, SH-SY5Y cells were seeded in 96-well plates at a density of 3.0 × 10⁴ cells per well and allowed to adhere for 24 h. Treatments were initiated when cells reached about 40–50% confluency. Cells were then incubated with HEWL assemblies at final concentrations of 1 or 7 μM for 48 h. Vehicle controls were prepared by diluting the corresponding buffer in medium at the same ratio (1:9, v/v) to ensure identical buffer exposure across all conditions.

### 4.12. Cell Viability and Cytotoxicity Assays

Cell viability was assessed using the Cell Counting Kit-8 (CCK-8; Dojindo, Japan), and absorbance was measured at 450 nm using a microplate reader. The CCK-8 results are expressed as the relative viability normalized to the vehicle control. Cytotoxicity was quantified by measuring lactate dehydrogenase (LDH) release using the CytoTox 96 Non-Radioactive Cytotoxicity Assay (Promega, USA), and absorbance was measured at 490 nm. LDH release was normalized to that of a maximum release lysis control prepared according to the manufacturer’s protocol.

### 4.13. Live/Dead Assay

Cell viability was evaluated using a live/dead fluorescence assay (Live/Dead Viability/Cytotoxicity Kit, L3224; Thermo Fisher Scientific). After 1 h of treatment with the HEWL assemblies, the cells were washed with PBS and incubated with the staining solution for 15–20 min at room temperature in the dark. Fluorescence images were acquired using a fluorescence microscope under identical acquisition settings across all experimental groups. Live (calcein-AM, green) and dead (ethidium homodimer-1, red) cells were quantified and reported as percentages of the total cells.

### 4.14. Immunofluorescence Staining

For fluorescence imaging, SH-SY5Y cells were cultured in 8-well chamber slides. Cells were seeded at 3.0 × 10⁵ cells mL⁻¹ (300 μL per well) and allowed to attach. HEWL assemblies were added to the cells and fixed immediately. The cells were fixed with 3.7% paraformaldehyde in PBS for 10 min and permeabilized with 0.1% Triton X-100. F-actin was stained using Alexa Fluor 568 phalloidin (Thermo Fisher Scientific; A12380) according to the manufacturer’s instructions, and the nuclei were counterstained with 4′,6-diamidino-2-phenylindole (DAPI). The samples were mounted using the VECTASHIELD Antifade Mounting Medium (Vector Laboratories). Fluorescent images were acquired using a confocal laser scanning microscope (LSM700, Carl Zeiss, Germany).

### 4.15. Image Acquisition and Quantification

Fluorescence images were acquired under identical microscopic settings for quantitative comparison. F-actin organization was quantified in Fiji (ImageJ, National Institutes of Health, USA) using a consistent threshold across conditions and calculating the F-actin-positive area fraction (thresholded area/total image area). Nuclear morphology was analyzed from DAPI images using StarDist 2D segmentation in Fiji; segmented nuclei were exported to the ROI Manager and circularity was computed as *4πA/P²*, where *A* is nuclear area and *P* is perimeter [56].

### 4.16. Statistical Analysis

Statistical analyses were performed using GraphPad Prism software (GraphPad Software, USA). Multiple group comparisons were conducted using one-way analysis of variance (ANOVA) followed by Tukey’s post hoc test. Data are presented as the mean ± SD, and *p* < 0.05 was considered statistically significant.

## AUTHOR INFORMATION

### Corresponding Author

Gyudo Lee

### Author Contributions

S.R. and E.S., contributed equally to this work.

Seokbeom Roh: Conceptualization, Investigation, Data curation, Methodology, Formal analysis, Visualization, Writing – original draft.

Eunji Song: Investigation, Data curation, Methodology, Formal analysis, Writing – original draft.

Minwoo Bae: Investigation, Data curation, Methodology, Validation

Da Yeon Cheong: Investigation, Methodology, Validation.

Sechan Han: Investigation, Validation.

Taeha Lee: Investigation, Validation.

Dain Kang: Investigation, Formal analysis. Hyungbeen Lee: Investigation, Formal analysis. Insu Park: Investigation, Methodology

Gyudo Lee: Conceptualization, Supervision, Funding acquisition, Project administration, Writing – original draft.

## ACKNOWLEDGMENTS

The authors acknowledge Mr. Jongwon Mun and Prof. Hyeongjin Lee for helpful assistance with confocal microscopy experiments related to cytoskeletal imaging.

## ABBREVIATIONS

ANS: 8-anilinonaphthalene-1-sulfonic acid;
AWI: air–water interface;
AFM: atomic force microscopy;
CCK-8: cell counting kit-8;
CD: circular dichroism;
DAPI: 4′,6-diamidino-2-phenylindole;
*H_AWI_*: air–water interface height; chemokine receptor 5;
HEWL: hen egg-white lysozyme;
LDH: lactate dehydrogenase;
PBS: phosphate-buffered saline;
PF-QNM: quantitative peak force nanomechanical mapping;
ThT: thioflavin T.

