## Supplementary Information for "Mechanochemically Programmed, Oligomer-Selective Amyloid Assembly via Axial Rotation"

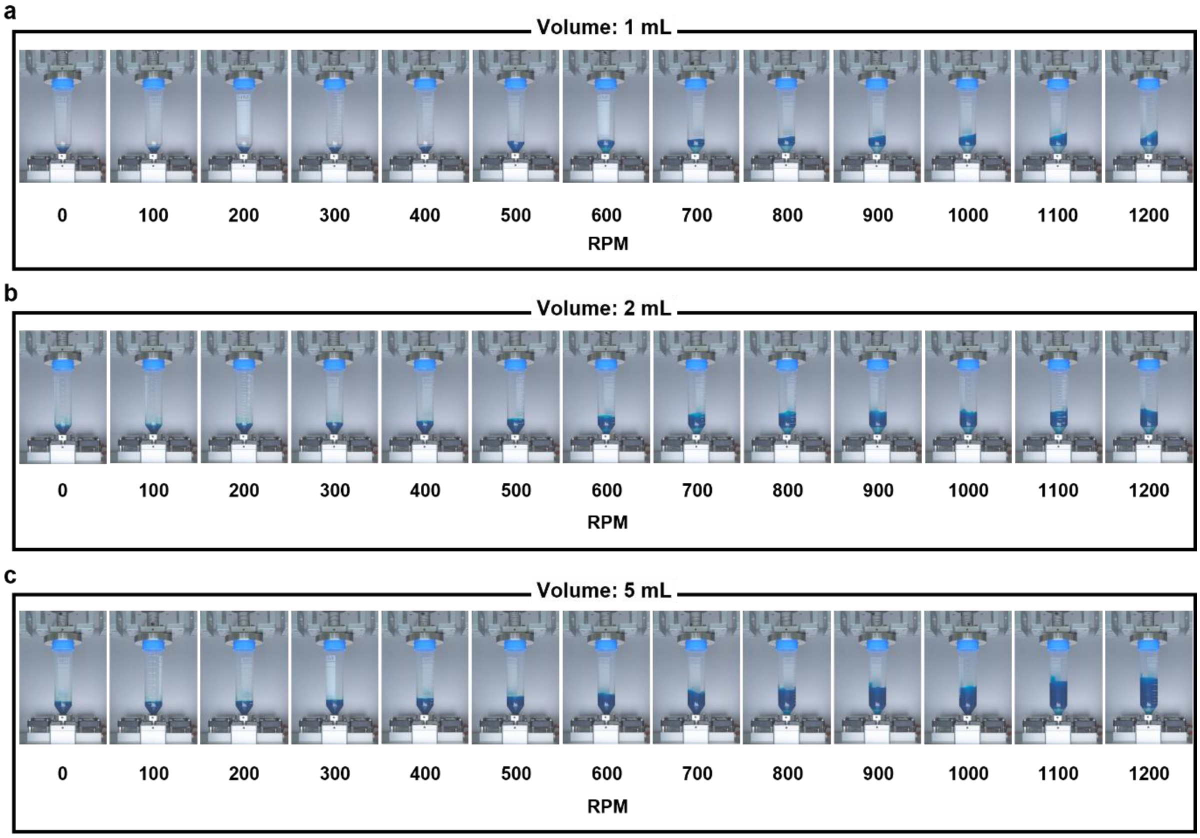


**Figure S1. Volume- and RPM-dependent deformation of the air–water interface.**

Representative images of the liquid interface at (a) 1 mL, (b) 2 mL, (c) 5 mL across increasing rotational speeds. Progressive wall climbing and interfacial tilting are observed with increasing RPM.

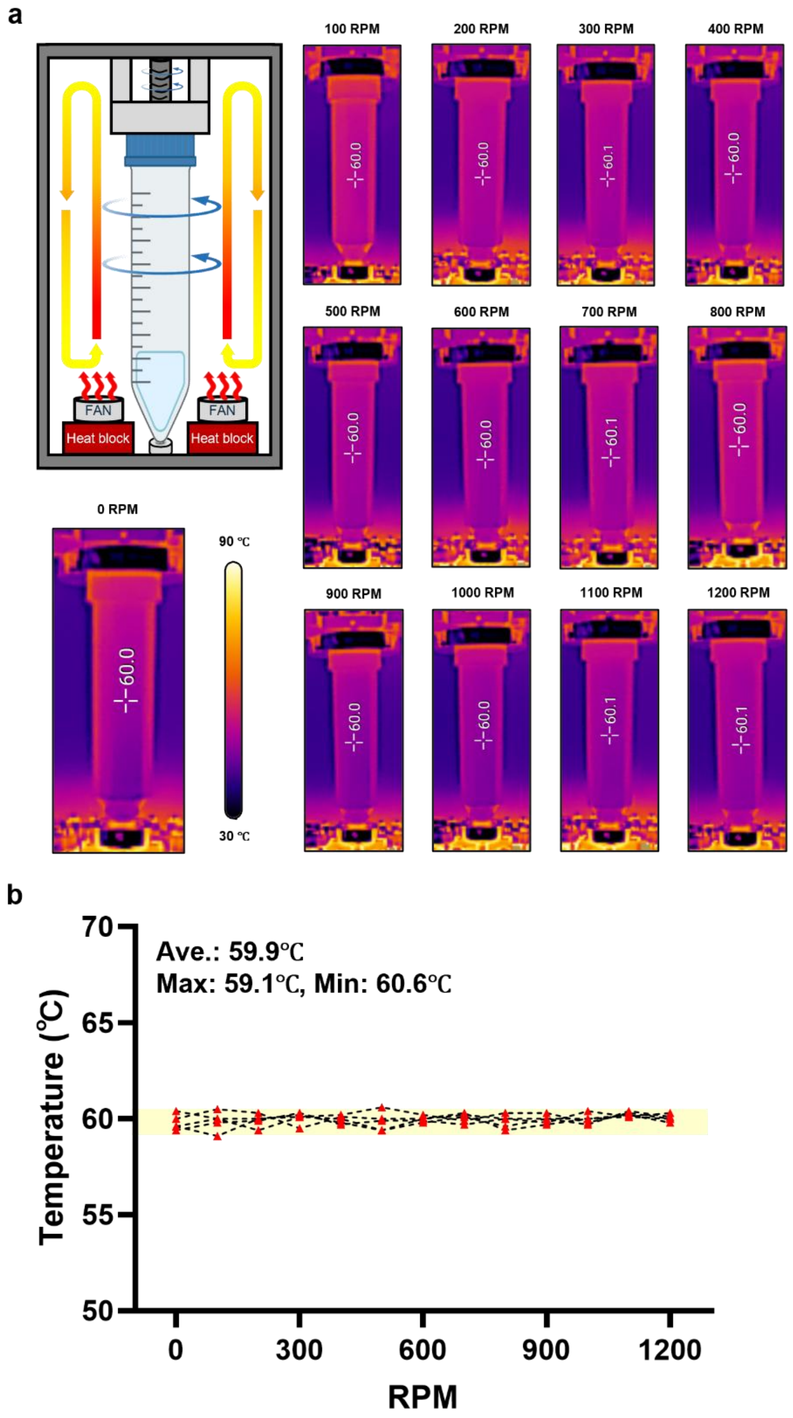


**Figure S2. Surface temperature stability of the rotating tube across RPM conditions.**

(a) Schematic of the axial rotation system and representative infrared thermal image used for surface temperature analysis. The conical tube was mounted on the axial rotator positioned in front of heat blocks, and thermal distribution was monitored using an infrared thermal imaging camera. The thermal map shows the spatial temperature profile of the tube surface under rotational conditions, with the color scale indicating temperature (30–90°C).(b) Quantitative analysis of tube surface temperature as a function of rotational speed (0–1200 RPM). Temperature values were extracted from a defined region of interest along the central tube wall in the infrared thermal images. Data are presented as mean ± SD (n = 4).

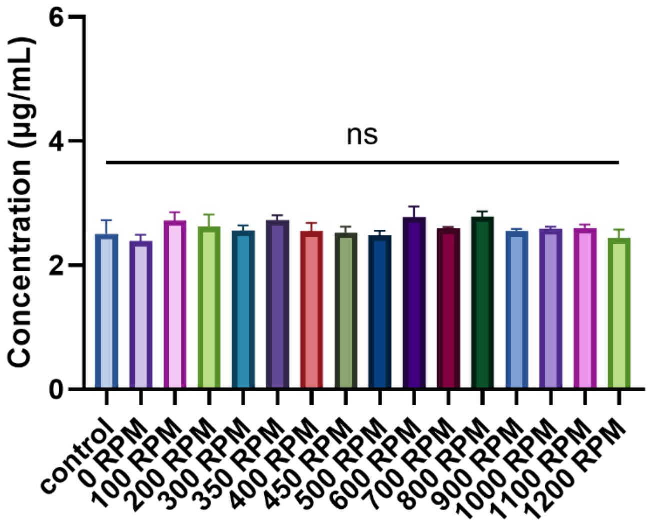


**Figure S3. Protein concentration analysis by bicinchoninic acid (BCA) assay.**

Protein concentration of samples prepared under control and various rotational speeds (0–1200 RPM) measured using the BCA assay. Concentrations are expressed in µg/mL. Data are presented as mean ± SD (n = 3). No statistically significant differences were observed among conditions (ns).

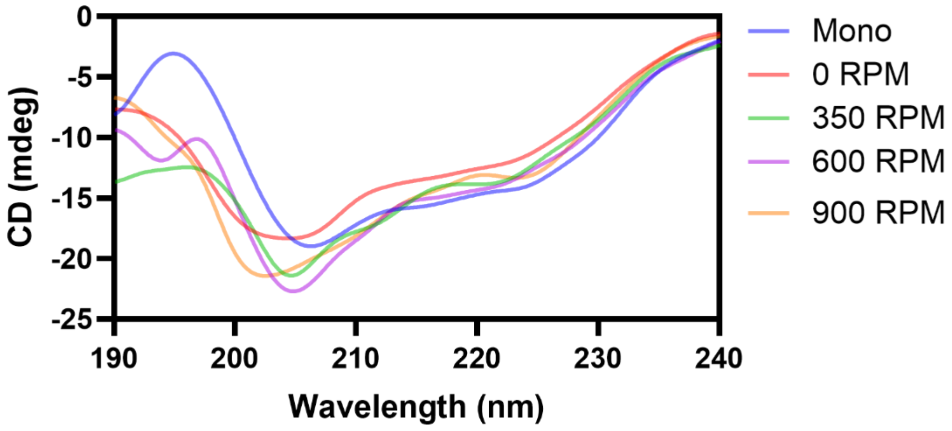


**Figure S4. Circular dichroism (CD) spectra of amyloid samples prepared under different rotational conditions.**

Far-UV CD spectra (190–240 nm) of monomer (Mono) and samples generated at 0, 350, 600, and 900 RPM. All spectra were acquired under PBS buffer and instrumental conditions.

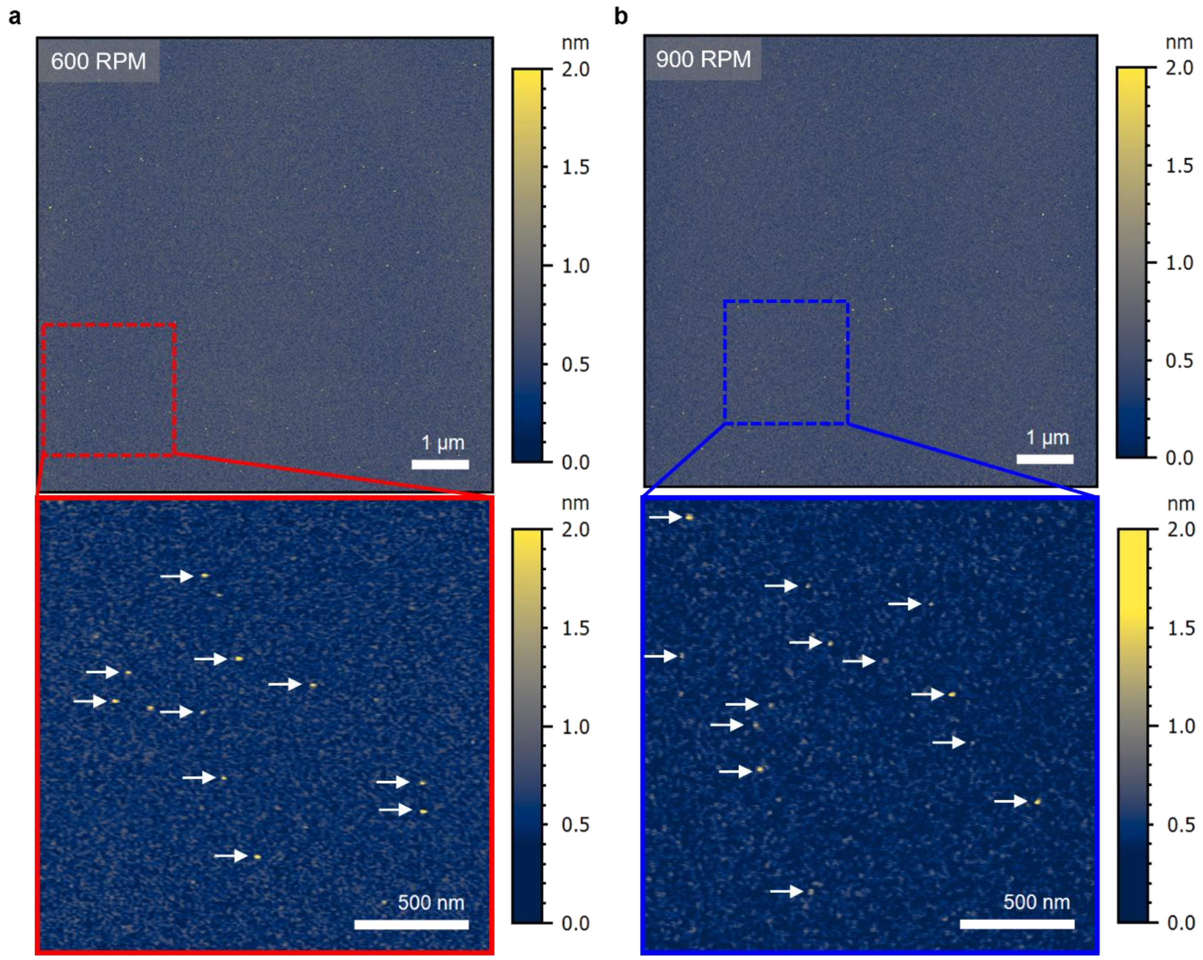


**Figure S5. High-resolution AFM analysis of oligomeric species formed at high rotational speeds.**

Representative atomic force microscopy (AFM) height images of samples prepared at (a) 600 RPM and (b) 900 RPM after incubation at 60 °C for 3 days. Dashed boxes indicate regions selected for magnified views shown below. White arrows denote representative oligomers.

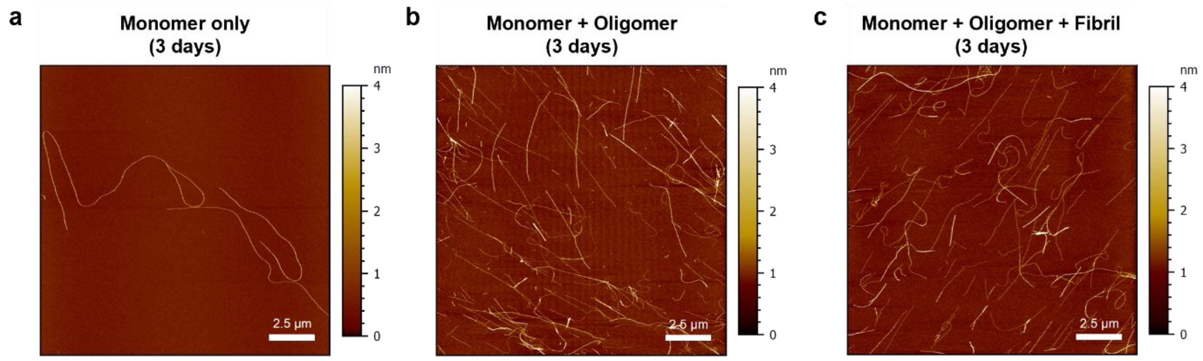


**Figure S6. AFM characterization of aggregates formed in the seeding experiments (corresponding to Figure 4).**

Representative atomic force microscopy (AFM) height images of samples collected after 3 days of incubation under three conditions: (a) monomer only, (b) monomer mixed 1:1 with oligomer-dominant assemblies generated at 600 RPM, and (c) monomer mixed 1:1 with oligomer- and fibril-containing assemblies generated at 350 RPM.

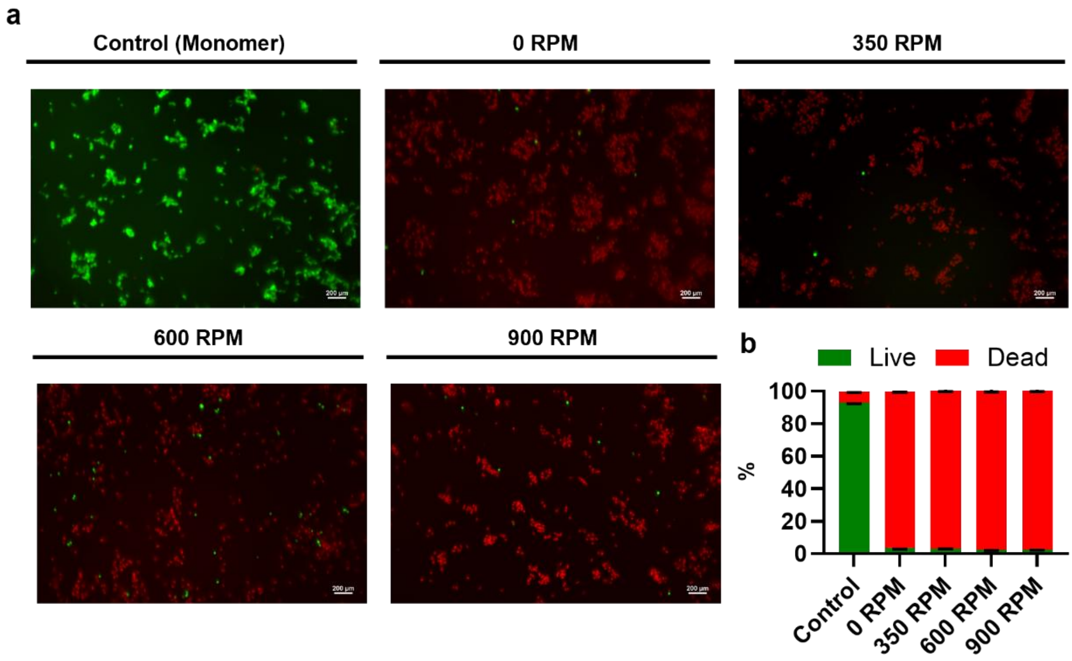


**Figure S7. Live/dead assay of SH-SY5Y cells treated with RPM-defined HEWL assemblies at 7 μM.**

HEWL samples were prepared at 60 °C for 3 days under different rotational speeds (Monomer, 0, 350, 600, and 900 RPM) and applied to SH-SY5Y cells at 7 μM. (a) Representative fluorescence images of live/dead staining. Live cells are shown in green and dead cells in red. Scale bars: 200 μm.(b) Quantification of live and dead cell populations expressed as percentage of total cells for each condition.

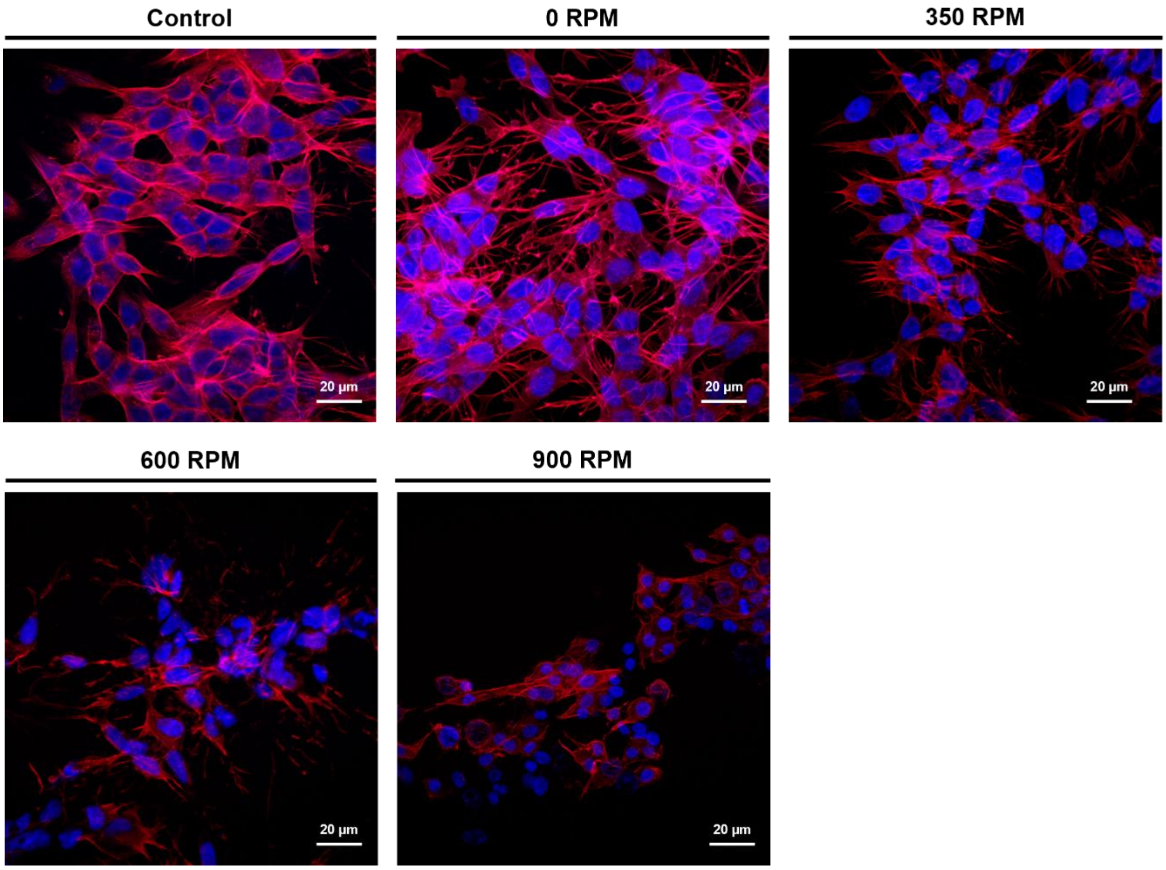


**Figure S8. High-magnification fluorescence images of cytoskeletal and nuclear staining (corresponding to Figure 6).**

Representative confocal fluorescence images of SH-SY5Y cells treated with RPM-defined HEWL assemblies (Monomer, 0, 350, 600, and 900 RPM; 60 °C, 3 days). F-actin was stained with Phalloidin–TRITC (red), and nuclei were stained with DAPI (blue). Merged images are shown. Scale bars: 20 µm.
